# Comprehensive analysis of macrophages infected with avirulent and virulent *Mycobacterium tuberculosis* uncovers distinct immune mechanisms and anti-TB effect of treated exosomes

**DOI:** 10.1101/2024.12.24.630227

**Authors:** Li Yang, Lingna Lyu, Cuidan Li, Xiuli Zhang, Yingjiao Ju, Ju Zhang, Jie Liu, Liya Yue, Xiangli Zhang, Dandan Lu, Tingting Yang, Peihan Wang, Jie Wang, Xiaotong Wang, Sihong Xu, Yongjie Sheng, Chunlai Jiang, Jing Wang, Xin Hu, Tuohetaerbaike Bahetibieke, Zongde Zhang, Fei Chen

**Author notes:** To whom correspondence should be addressed to Fei Chen. Correspondence may also be addressed to Zongde Zhang. These authors contributed equally to this work (Li Yang, Lingna Lyu, Cuidan Li, Xiuli Zhang, Yingjiao Ju, Ju Zhang).

## Abstract

Tuberculosis (TB) is now the world’s second deadliest infectious killer after COVID-19. Human-macrophages and their secreted exosomes play important roles in combating invading *Mtb*. However, the panoramic analysis of the underlying immune mechanism for the infected macrophages, package or secretion mechanism, and anti-TB effect of *Mtb* treated exosomes remain poorly understood. Here we conducted comprehensive analyses of the macrophages infected with avirulent and virulent *Mtb* (H37Ra & H37Rv) and their secreted exosomes, collected cells and corresponding exosomes for omics and phenotypic analysis. The results showed that avirulent *Mtb* stimulated robust immune-responses and apoptosis in macrophages to eliminate the invading *Mtb*; virulent *Mtb* induced severe necrosis and immune-escape for survival. The HMGB1 signaling pathway and *TNFRSF1B* plays important roles in the immune escape of virulent *Mtb*. Interestingly, our results suggest that macrophages kill *Mtb* in an IFN-γ independent but simulative way, highlighting the central role of IFN signaling pathway in anti-TB immune response. Moreover, we observed selective transport of host and *Mtb* RNAs from macrophages to exosomes. Notably, “H37Ra-treated exosomes” displayed a higher anti-TB effect than “H37Rv-treated exosomes” due to some enriched pro-inflammation and immune-escape related *Mtb* proteins in these two exosomes, respectively. Conclusively, our findings shed new light on the immune mechanism of macrophages in response to *Mtb* infection, offering a new TB-treatment strategy and some promising vaccine candidates.

## Introduction

Tuberculosis (TB) has been the leading lethal infectious disease worldwide from a single infectious agent, *Mycobacterium tuberculosis* (*Mtb*), for a long time (2014 – 2019) and is now the world’s second deadliest infectious killer (2020 – 2022) after COVID-19 [1–5]. According to the WHO Global TB report 2022, an estimated 10.6 million people were diagnosed with TB in 2021, a 4.5% increase from 2020, reversing declines of ∼2% per year for the past two decades. Similarly, 1.6 million people died from TB in 2021, reversing 14 years of decline from 2005 to 2019 [3].

It has been reported that immune mechanism involved various immune cells are quite complex after *Mtb* infection [6–8]. Among them, macrophage is the most critical one to protect host against *Mtb* through triggering off multiple innate immune responses (including recognition, phagocytosis, and lysis of intracellular *Mtb*) [8]. Further, macrophages can stimulate cascading acquired immune responses to kill intracellular *Mtb* (including antigens presentation, activation of specific CD4+T cells, production of IFN-γ and other important antibacterial cytokines, and macrophage activation for phagocytosing/killing the invading *Mtb*) [9]. Previous studies have also shown that several signaling pathways play vital roles in immune responses in macrophages, such as PRRs represented by Toll-like receptor (TLR), autophagic signaling pathway, and interferon (IFN) signaling pathway [8].

Further researches explored different infection and immune mechanisms of the virulent H37Rv and avirulent H37Ra *Mtb* reference strains on macrophages. Zhang *et al.* reported a faster growth rate of H37Rv than H37Ra in human monocyte-derived macrophages (hMDMs) [10]; Chen *et al.* found that H37Rv, but not H37Ra, could cause necrosis by significant disruption of the mitochondrial inner membrane in hMDMs [11]; Freeman *et al.* demonstrated differential growth and cytokine/chemokine induction in H37Ra and H37Rv infected murine macrophages [12]. There are also several transcriptomic studies for the macrophages infected with H37Rv and H37Ra: Kalam *et al.* reported extensive remodeling of alternate splicing in the infected macrophage [13]; Lee *et al* identified many differentially expressed genes (DEGs) related to innate immune responses in H37Ra and H37Rv infected hMDMs, and further confirmed the important role of *slc7a2* (solid carrier family 7 member 2) in *Mtb* survival [14]; Pu *et al.* found that H37Rv infection could trigger off more severe inflammatory immune responses for facilitating tissue damage than H37Ra and BCG infections, through comparative transcriptomic analysis of infected THP-1-derived macrophages [15].

In recent years, exosomes have been reported to play important roles in various diseases, including TB [16]. Exosomes are extracellular, cup-shaped membrane vesicle (∼100 nm) that mediate signal transmission and substance transferring between cells by trafficking bio-active molecules (including DNA, RNA, protein and lipid) [17]. Secreted exosomes from infected cells are closely related to innate and acquired immune responses of the body through packing *Mtb* and human contents (including protein and RNAs) [18, 19]. Giri *et al.* identified 41 highly antigenic and bio-functional *Mtb* proteins in the exosomes from the J774 cells infected by H37Rv using LC-MS/MS method [20]; Obregón-Henao *et al.* found that the *Mtb* RNAs in exosomes could induce early apoptosis in human monocytes [21]. Our previous studies also showed many human and *Mtb* RNAs in serum exosomes from latent and active tuberculosis, including mRNA, miRNA, and other small RNA, plenty of which were closely related to immune responses [22, 23].

Overall, human immune responses in various immune cells decide the outcome of *Mtb* infection, among which macrophages play a crucial role in fighting against invading *Mtb*. However, a panoramic analysis of the immune mechanism in infected macrophages is still lacking. The second concern is to explore differential immune mechanisms of human macrophages in response to the virulent H37Rv and avirulent H37Ra reference strains, and find out why H37Ra can be eliminated by macrophages while H37Rv survives in *vivo* and in *vitro*. The third concern is to elucidate the packaging or secretion mechanism of host and *Mtb* RNAs into exosomes from infected macrophages, and explore whether the *Mtb* treated exosomes help macrophages resist *Mtb* infection and which genes in exosomes may contribute to the antibacterial effect.

## Results

### Differential survival outcomes of macrophages and invading *Mtb* between avirulent and virulent *Mtb* infections

To explore the infection mechanism of *Mtb* on human host cells, we infected PMA-differentiated THP-1 macrophages with the virulent H37Rv and avirulent H37Ra reference strains (Figure 1), and confirmed the infection of both strains through Ziehl-Neelsen staining (Figure 2A). Further, Methyl thiazolyl tetrazolium (MTT) and colony-forming unit (CFU) assays were conducted to measure cell growth and the invasion of H37Ra or H37Rv, respectively (Figure 2B, C). It showed different cell viabilities between the macrophages infected by H37Rv and H37Ra. The growth of H37Rv-infected macrophages increased during the first 24 hours of infection and significantly declined thereafter (similar as controls), whereas the growth of H37Ra-infected macrophages continuously declined from the beginning (Figure 2B). Correspondingly, the CFU assay revealed differential survival of invading H37Rv and H37Ra, with the bacterial load of H37Rv increasing during the first 24 hours of infection and gradually decreasing thereafter, while that of H37Ra showed a continuous decrease from the beginning of infection (Figure 2C). Here, we washed off uninfected *Mtb* from the cells to ensure the reliability of the CFU assay for invading *Mtb*.

**Figure 1.**
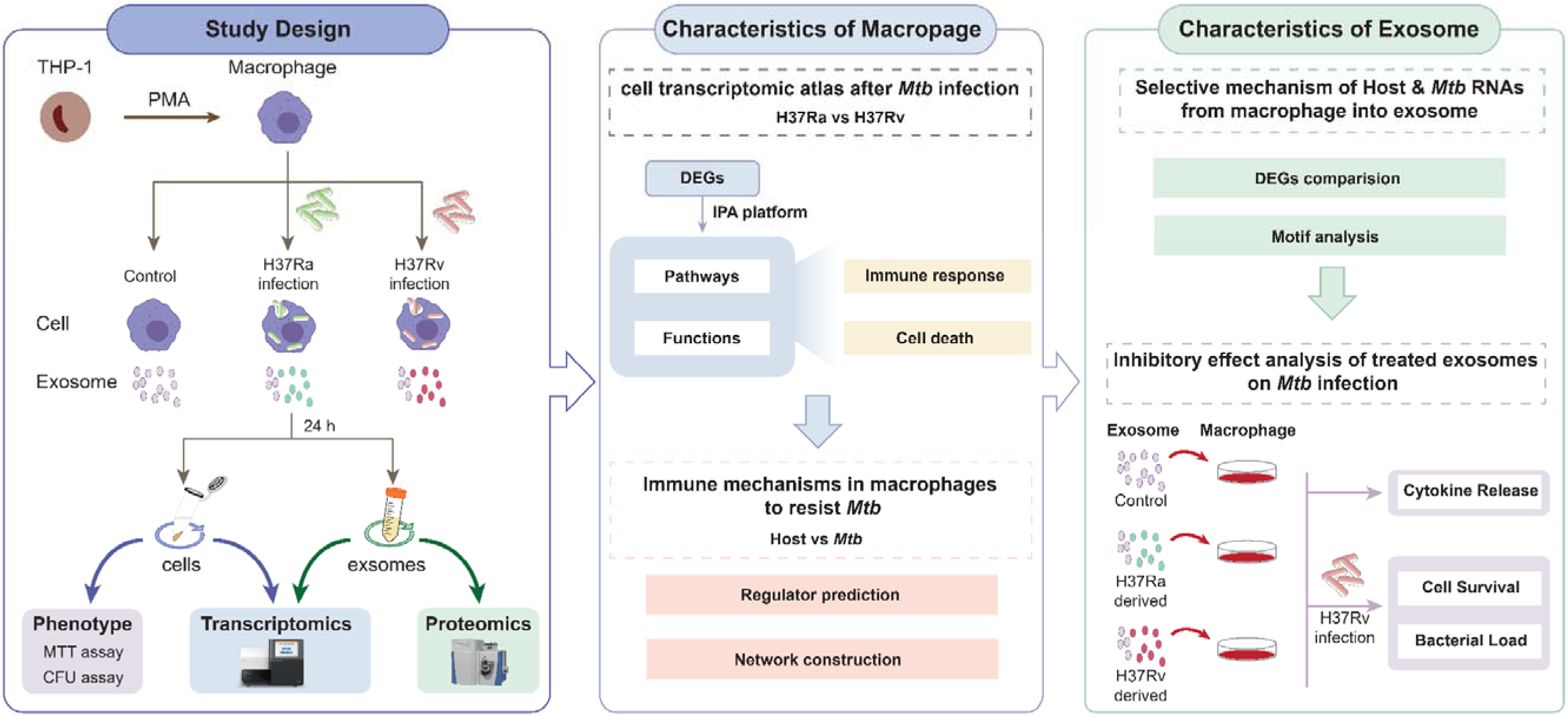
Diagram showing the schematic workflow including bioinformatic data analysis and experiments of this study. (1) Virulent and avirulent H37Ra reference strains were used to infect PMA-differentiated THP-1 macrophages. Cells and corresponding exosomes ollected for phenotypes, transcriptomics and proteomics study (left panel); (2) Characteristics of infected macrophages were further ed, including the transcriptomic atlas after *Mtb* infection and the immune mechanism in macrophages to resist *Mtb* (middle panel); (3) teristics of secreted exosomes were then analyzed, including the selective mechanisms and the anti-TB effect of the treated exosomes panel).

**Figure 2.**
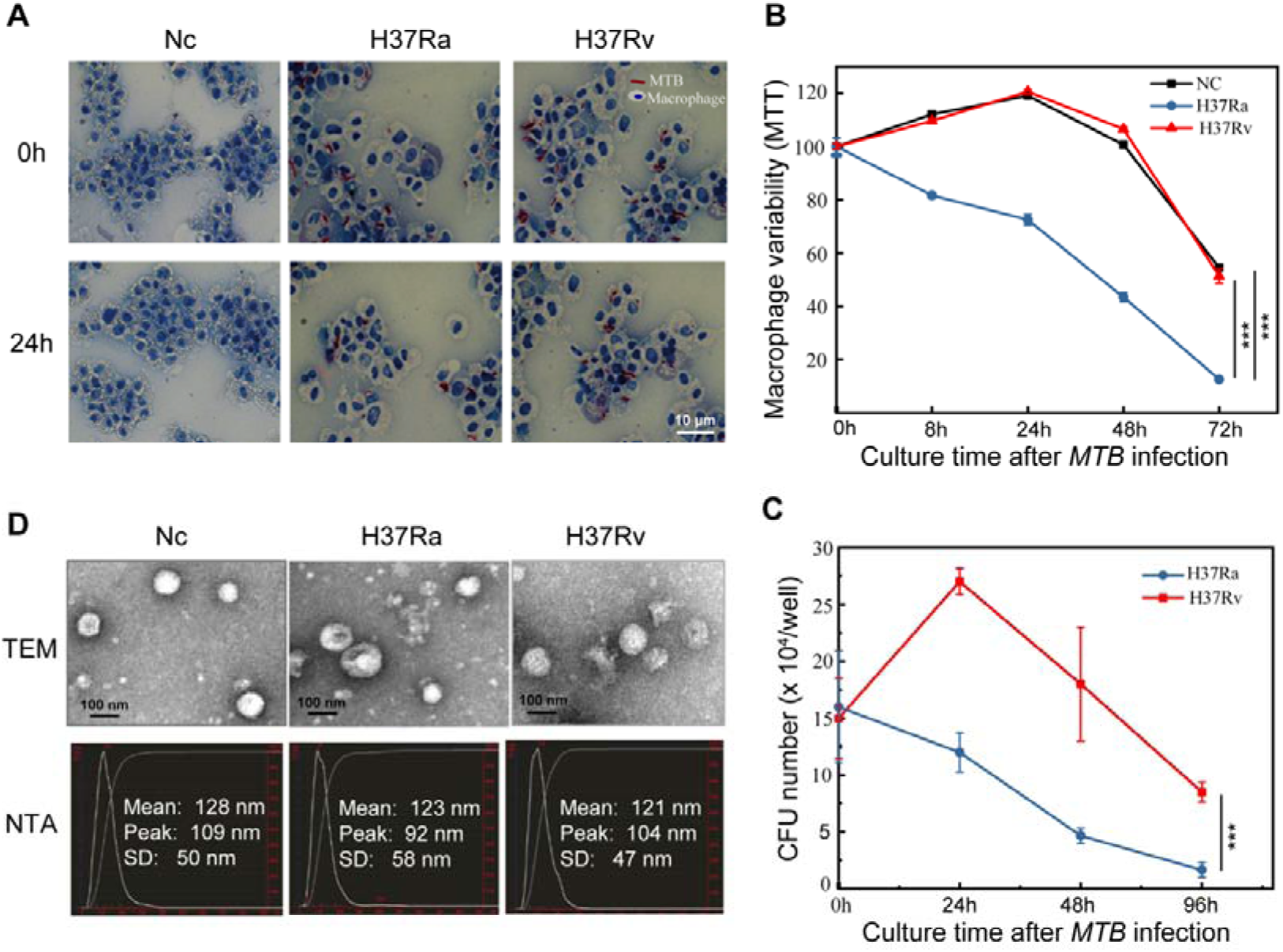
Differentially survival status of macrophages and invading *Mtb* between H37Ra and H37Rv infections. **(A)** Ziehl-Neelsen staining shows efficient *Mtb* infections in macrophages (MOI = 10) after 24 h infection. *Mtb* and cells are stained red and blue, respectively. **(B)** MTT assay showing differentially viability of macrophages infected with H37Ra and H37Rv (MOI = 10). The absorbance values are normalized to the uninfected control cells. **(C)** CFU assay showing the viability of the invading *Mtb* in macrophages infected with H37Ra and H37Rv (MOI = 10). Data in (B) and (C) are from three independent experiments with biological duplicates in each (mean ± SEM of n = 3 duplicates). ***, p < 0.001. (D) Exosome morphology and size distribution characterizations by Transmission Electron Microscope (TEM) and Nanoparticle Tracking Analysis (NTA); The x-axis represents the size (diameter in nm), and the y-axis represents the concentration (number of particles per ml). The red color highlights the scales of the x and y axes, which we have added to the figure. Additionally, this figure shows the three main values for the diameter of exosomes: the mean, peak, and standard deviation (SD) (peak: ∼ 100 nm).

The similar growth curves for the survival macrophages and invading *Mtb* after infection indicated a correspondence between them, which can also be reasonably explained by previous studies [24, 25]: for macrophages infected with H37Rv, immune escape of virulent *Mtb* in the early infection stage helps them to survive in the host cells, followed by gradual elimination by increasing immune responses; in contrast, the avirulent H37Ra is continuously eliminated by activated immune system of macrophages from the beginning of infection.

### Significant transcriptomic changes in the macrophages and released exosomes caused by avirulent and virulent *Mtb* infections

After 24 hours of H37Rv and H37Ra infection, both macrophages and their released exosomes were collected for transcriptomic analysis. The purity and the integrity of the exosomes were confirmed using transmission electron microscopy (TEM) and nanoparticle tracking analysis (NTA) (Figure 2D). Total RNAs were then extracted from the macrophage cells and exosomes, which displayed characteristic cell and exosomal RNA profiles (Figure S1A) as previously reported [26]: the cellular RNAs contained typical eukaryotic 5S, 18S and 28S ribosomal RNAs, while the exosomal RNAs had a diffuse distribution with high abundance of short RNA fragments.

Transcriptomic sequencing was then conducted on infected macrophage cells and their secreted exosomes (Figure S1B, C). Compared with controls, both avirulent and virulent *Mtb* caused significant transcriptomic changes in the macrophages and released exosomes (Figure 3A, B): 208 and 650 differentially expressed genes (DEGs) were identified in the H37Rv infected cells and their released exosomes, respectively; 256 and 548 DEGs were identified in the H37Ra infected cells and their released exosomes, respectively (fold change ≥ 2, p-value ≤ 0.05).

**Figure 3.**
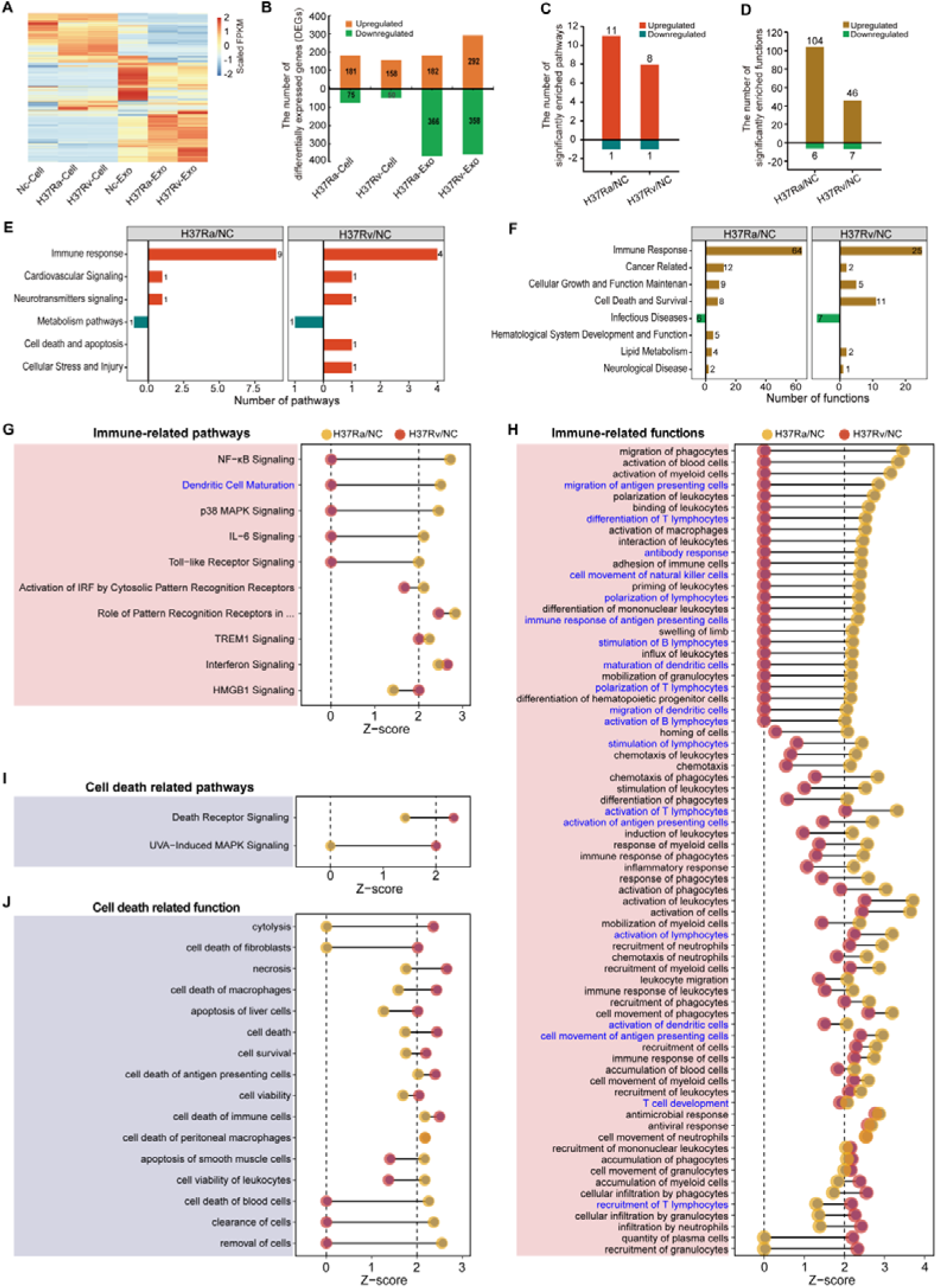
Expression profiles and significantly enriched pathways and biofunctions of the *Mtb* infected macrophages and released exosomes. **(A)** Heatmap showing the differentially expressed genes among the six groups (|Fold change| ≥ 2, p < 0.05, q < 0.05). **(B)** Bar plot showing significantly up-/down-regulated mRNAs in the *Mtb* infected cells and released exosomes. Red and blue bars represent up-regulated and down-regulated mRNAs, respectively (|Fold change| ≥ 2, p < 0.05). **(C, D)** Bar plots showing significantly enriched pathways and biofunctions (D) in the H37Rv/NC and H37Ra/NC groups. **(E, F)** Bar plots showing the significantly enriched pathways (E) / biofunctions (F) in six / eight aspects in the H37Rv/NC and H37Ra/NC groups. Red and yellow bars indicate the significantly upregulated/activated pathways and biofunctions (Z-score ≥ 2, p < 0.05), respectively; blue and green bars represent the significantly inhibited/downregulated pathways and biofunctions (Z-score ≤ −2, p < 0.05), respectively. **(G, H)** Dumbbell charts showing the innate and adaptive immunity pathways (G) and biofunctions (H) between the two groups. Blue font represents adaptive immunity pathways and biofunctions. **(I, J)** Dumbbell charts showing the cell apoptosis/death related pathways (I) and functions (J) between the two groups.

### Avirulent *Mtb* causes stronger immune responses and apoptosis, while virulent *Mtb* causes stronger immune escape and non-apoptotic programmed cell death in macrophages

To explore the immune mechanism of human macrophages in response to *Mtb* infection, we analyzed significantly enriched pathways and biofunctions using IPA (Ingenuity Pathway Analysis). Compared to controls, most of the enriched pathways and biofunctions were significantly upregulated (p value < 0.05, Z-score ≥ 2) in the Ra/NC and Rv/NC groups: 11 (11/12, 91.67%) and eight (8/9, 88.89%) pathways were significantly upregulated in the Ra/NC and Rv/NC groups, respectively, and only one pathway (PPAR signaling) was significantly downregulated in both groups (Figure 3C); 104 (104/110, 94.55%) and 46 (46/53, 86.79%) biofunctions were significantly upregulated, and only six and seven significantly downregulated ones in the Ra/NC and Rv/NC groups, respectively (Figure 3D). In addition, we also observed that the Ra/NC group exhibited a total of 12 significantly upregulated pathways and 110 activated biofunctions, respectively, both of which were higher compared to the corresponding numbers in the Rv/NC group.

These significantly enriched pathways and biofunctions were then classified into six and eight aspects, respectively, based on their characteristics (Figure 3E, F). Notably, the immune-response related pathways and biofunctions ranked first among all aspects and were significantly upregulated in both Ra/NC and Rv/NC groups. This demonstrates that human macrophages resist *Mtb* infections through activating multiple immune-response related pathways and functions.

Further, there were higher activation Z-score for immune-response related pathways and functions in the H37Ra infected macrophages than in the H37Rv macrophages, suggesting a stronger immune response of macrophages against avirulent H37Ra infection and immune escape of macrophages for H37Rv infection (Figure 3E-H), which was in agreement with their distinct phenotypic outcomes (Figure 2B, C). Among all 10 significantly activated immune-related pathways in the Ra/NC and Rv/NC groups (Figure 4G), six were specifically upregulated in the Ra/NC group, including four innate immune response related pathways (Toll-like receptor signaling, NF-κB signaling, *etc.*) and two acquired immune response related pathways (Dendritic cell maturation and IL-6 signaling). Among the three shared significantly activated pathways, two associated with pathogen recognition and immune response amplification were more activated in the Ra/NC group than in the Rv/NC group [“Role of Pattern Recognition Receptors in Recognition of Bacteria and Viruses” (Z-score: 2.828 vs. 2.449) and “TREM1 Signaling” (Z-score: 2.236 vs. 2)]. It is worth noting that the “Interferon Signaling” pathway was almost identically activated in both H37Ra and H37Rv infected macrophages (Z-score: 2.449 vs. 2.646). Specifically, only one immune related pathway “HMGB1 signaling” was specifically activated in the H37Rv-infected macrophage among all the 10 pathways.

**Figure 4.**
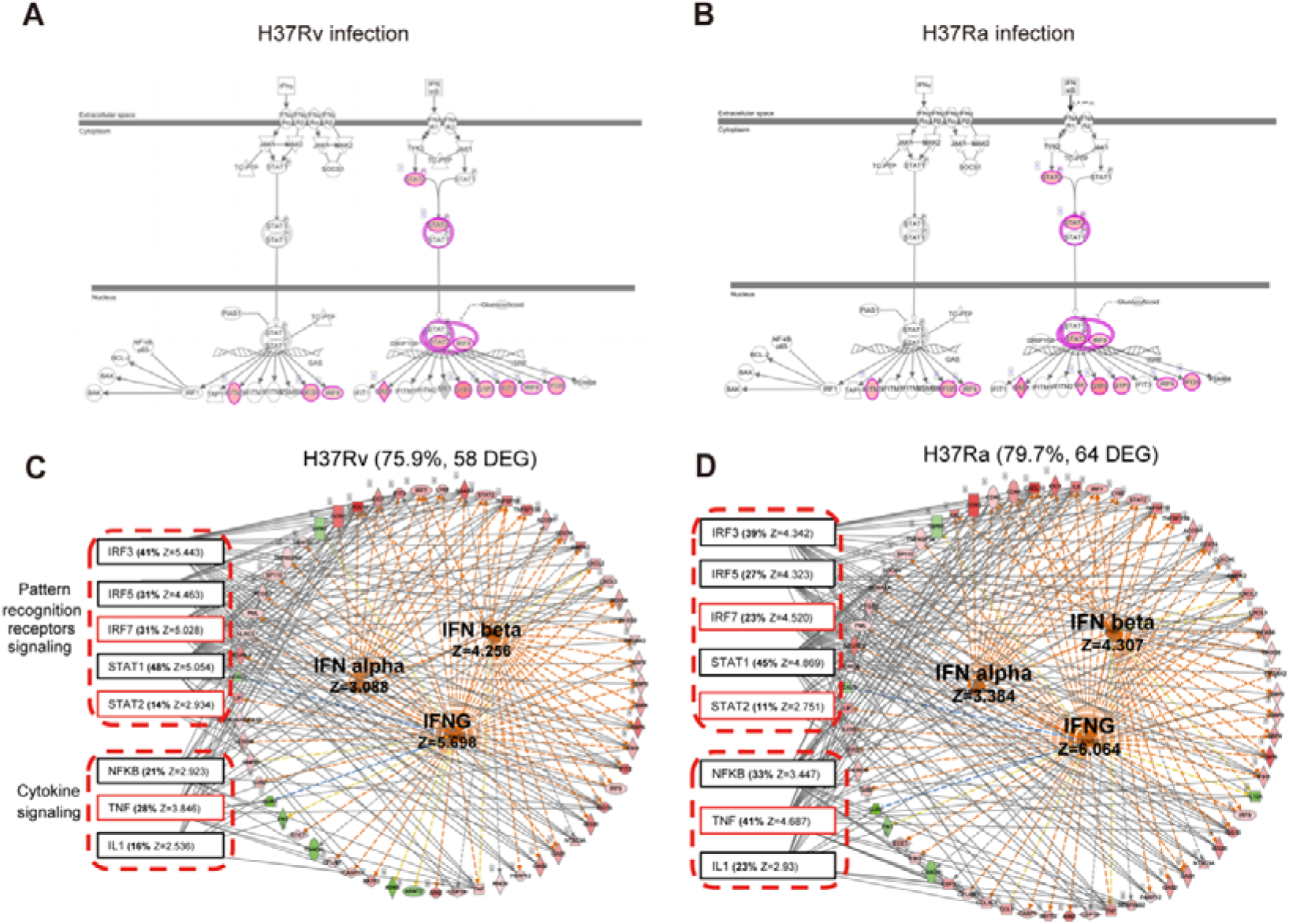
Simulative activation of interferon signaling pathways in macrophages through activating the downstream genes of interferons. **(A, B)** The activation of interferon signaling pathway after H37Ra and H37Rv infection. Pink circles represent differential genes and filled colors represent fold change. **(C, D)** Interferon pathway simulation network of H37Ra and H37Rv infection. The IFNs in the circle are predicted upstream regulators and the circular arrangement of genes on the right represents the downstream DEGs that support interferon activation. The left eight predicted upstream regulators in dashed boxes are predicted to be the target molecules in simulation networks which have common downstream DEGs interferon activation.

On the other hand, among all 72 enriched immune-response related biofunctions in the two groups (Figure 3H), a significant proportion (65.3%, 47/72) were specifically activated in the Ra/NC group, including 32 innate immune response functions related to phagocytosis, recruitment, migration, and chemotaxis (“migration of phagocytes”, “activation of myeloid cells”, *etc.*) and 15 acquired immune response functions concerning T/B lymphocytes activation (differentiation of T lymphocytes, activation of B lymphocytes). The majority of shared up-regulated functions (14/17, 82.35%) were also more activated in the Ra/NC group than in the Rv/NC group, including 11 innate immune-related functions (“activation of leukocytes”, “activation of lymphocytes”, *etc.*) and three acquired immune-related ones (“activation of T lymphocytes”, “activation of lymphocytes”, *etc.*).

In addition, we observed more significantly activated non-apoptotic programmed cell death-related pathways and biofunctions in H37Rv infected macrophages than in H37Ra infected macrophages (Figure 3I, J). On the one hand, both enriched pathways associated with cell deterioration/necrosis were specifically upregulated in the Rv/NC group (Figure 3I). On the other hand, among all the 16 enriched cell death-related functions in the two groups (Figure 3J), eight were specifically activated in the Rv/NC group, including “cytolysis”, “cell death of fibroblasts”, “necrosis”, and “cell death”, while two shared functions showed higher activation in the Rv/NC group than in the Ra/NC group (“cell death of antigen presenting cells” and “cell death of immune cells”). These results suggest more severe non-apoptotic programmed cell death in the H37Rv infected macrophages. Conversely, one apoptotic programmed cell death-related function was specifically activated in H37Ra infected macrophages (apoptosis of smooth muscle cells).

We then analyzed the top 10 upregulated DEGs in the H37Ra and H37Rv infected macrophages, respectively (Table 1), and found that eight of these genes were shared between both groups. Of these shared DEGs, five were related to immune response, including three cytokines (*CXCL8*, *CCL3*, and *CCL3L3*) and two innate immune response related genes (*IFI6* and *BST2*); four of these immune-related DEGs (including the three cytokines and *IFI6*) were showed higher expression in the H37Ra infected macrophages. In addition, among the four specific DEGs, two specifically upregulated DEGs (*FTL* and *FTH1*) in the avirulent H37Ra infected macrophages have been reported to be associated with production of reactive oxygen species for the clearance of pathogenic bacteria [27]; while one specifically upregulated DEG (*TDO2*) in the virulent H37Rv infected macrophages has been shown to be related to evade immune surveillance [28].

**Table 1.** The top10 differentially expressed genes in the H37Ra and H37Rv infected macrophages.

| Gene name | FPKM (H37Ra/NC) | Fold Change (H37Ra/NC) | Top10 (H37Ra/NC) | FPKM (H37Rv/NC) | Fold Change (H37Rv/NC) | Top10 (H37Rv/NC) | Function |
| --- | --- | --- | --- | --- | --- | --- | --- |
| *FTL* | 15757.2 | 2.094 | √ | - | - | - | Production of reactive oxygen species |
| *CXCL8* | 5729.65 | 4.453 | √ | 5193.77 | 4.037 | √ | Production of reactive oxygen species, Cell-To-Cell signaling and interaction, Cell Death and Survival |
| *FTH1* | 4884.62 | 2.036 | √ | - | - | - | Production of reactive oxygen species, Cell death and survival |
| *CCL3* | 3088.31 | 4.919 | √ | 1633.12 | 2.601 | √ | Cell death and survival, Cell-To-Cell signaling and interaction |
| *CCL3L3* | 2493.87 | 6.186 | √ | 1379.52 | 3.423 | √ | Cell-To-Cell signaling and interaction |
| *IFI6* | 2197.59 | 6.519 | √ | 2143.59 | 6.359 | √ | Innate immune response, Cell death and survival |
| *ISG15* | 940.247 | 22.608 | √ | 1275.98 | 30.681 | √ | Cell death and survival |
| *MFSD2A* | 729.472 | 3.689 | √ | 541.615 | 2.739 | √ | Cell-To-Cell signaling and interaction |
| *RGS1* | 631.227 | 2.647 | √ | 716.261 | 3.004 | √ | Cell-To-Cell signaling and interaction |
| *BST2* | 630.502 | 2.871 | √ | 718.411 | 3.272 | √ | Innate immune response |
| *TDO2* | - | - | - | 591.311 | 2.681 | √ | Cell Death and Survival |
| *LY6E* | - | - | - | 403.049 | 6.829 | √ | Cell-To-Cell Signaling and Interaction |

Note: The genes in red and blue represent the specifically enriched DEGs in the H37Ra and H37Rv infected macrophages, respectively.

Collectively, our findings from the enriched pathways/functions, top 10 DEGs and phenotypes (Figure 2, Figure 3, and Table 1) reveal the differential immune mechanisms of human macrophages in response to virulent H37Rv and avirulent H37Ra reference strains: H37Ra infection stimulates a more robust immune responses and apoptotic programmed cell death in macrophages to resist and clear invading *Mtb*, while H37Rv infection leads to more severe non-apoptotic programmed cell death and immune escape in macrophages for survival. This sheds light on why H37Ra can be eliminated by macrophages and H37Rv survived in macrophages.

### Simulative activation of interferon signaling pathways in macrophages against *Mtb* infection through activating the downstream genes of interferons

To further explore the immune mechanisms in macrophages to resist *Mtb*, we used IPA to predict the upstream regulators for all the DEGs. Surprisingly, among the top 25 predicted upstream regulators (Table S1, S2), three were classical interferons: IFN-γ belonging to type II interferon (Z = 6.194/5.698 for H37Ra/Rv infected cells), IFN-α belonging to type I interferon (Z = 5.244/4.758 for H37Ra/Rv infected cells) IFN-β1 belonging to type I interferon (Z = 4.307/4.256 for H37Ra/Rv infected cells). Notably, IFN-γ was ranked number one in all predicted upstream regulators for both H37Ra and H37Rv infected macrophages. However, it is well known that type II interferon IFN-γ is produced by activated CD4+T and NK cells rather than macrophage [29], which is also supported by our transcriptomic data where the expressions of all types of interferons were hardly detected in the *Mtb* infected macrophages (Table S3).

What caused the discrepancy between the predicted upstream regulators and the real transcriptomic data regarding interferons? We initially observed a significant activation of the “Interferon Signaling” pathways in both H37Ra and H37Rv infected macrophages (Figure 3G), which could be attributed to the upregulation of some downstream DEGs in the pathways, including seven DEGs (*STAT2*, *IRF9*, *OAS1*, *G1P2*, *G1P3*, *IFIT3*, *IFI135*) in Type I interferon signaling pathway and three DEGs (*IFITM3*, *IFI35*, *IRF9*) in Type II interferon signaling pathway (Figure 4A, B). Consequently, the three interferons were predicted to be significantly activated as upstream regulators in the “Interferon Signaling” pathways, although they were hardly expressed in the actual transcriptomic data.

Further analysis through constructing the upstream regulation network for interferon pathway revealed that many significantly activated downstream DEGs were further detected in all possible interferon signaling pathways (IFN-α, IFN-β, and IFN-γ), including 64 and 58 DEGs in H37Ra and H37Rv infected macrophages, respectively (Figure 4C, D). However, since the three interferons were not expressed in our *Mtb* infection model, we then searched for possible upstream regulatory genes to activate these downstream DEGs. Here, the top eight predicted upstream regulatory genes were responsible for the activation of most of these downstream DEGs, accounting for 79.7% (51/64) and 75.9% (44/58) of the H37Ra and H37Rv infected macrophages, respectively. Among these eight upstream genes, five were from “pattern recognition receptor related signaling” pathway and three were from “cytokine signaling” pathway.

Overall, although the interferons were hardly expressed in the transcriptomic data, the interferon signaling pathways/networks in macrophages could still be activated in a simulative manner to resist *Mtb* infection (Figure 3G, 4A, and 4B) through activation of some downstream genes (Figure 4C, D) by other upstream genes. Macrophages are able to kill *Mtb* in an IFN-γ independent but a simulative way, highlighting the critical role of the IFN signaling pathway in the anti-TB immune response of macrophages.

### Selective secretion of characteristic host RNAs from *Mtb* infected macrophages to exosomes

To uncover whether host RNAs are selectively secreted from *Mtb* infected macrophages into exosomes, we then compared the DEGs between the *Mtb* infected macrophages and their secreted exosomes (Figure 5A). The results showed a significantly difference in the RNA profiles between them, indicating selective secretion of some specific RNAs from *Mtb* infected macrophages into exosomes: only seven and six DEGs were shared in H37Rv or H37Ra infected cells and their exosomes, accounting for approximately 1% of all exosomal DEGs (7/650 for H37Rv infection, 6/548 for H37Ra infection); the abundances of all shared expressed genes (4,090/4,194 genes in H37Ra and H37Rv, FPKM ≥ 10) in the *Mtb* infected cells showed a poor correlation with their exosomes (Figure 5B).

**Figure 5.**
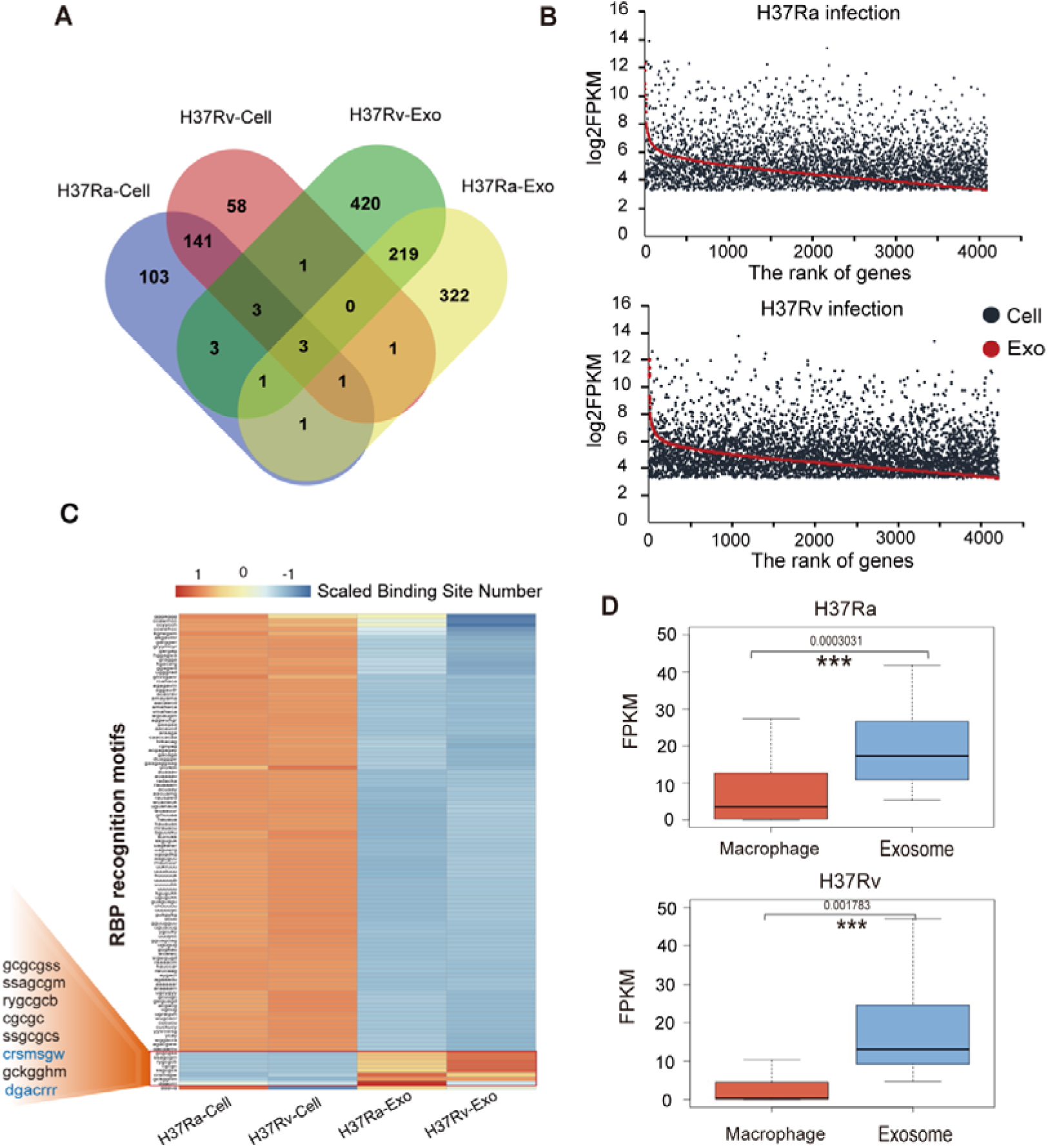
Selective secretion of characteristic host RNAs with RBP recognition motifs from *Mtb* infected macrophages to exosomes. **(A)** Venn diagram showing the shared genes among the *Mtb* infected cells and released exosomes (|Fold change| ≥ 2, p < 0.05, q < 0.05). **(B)** The abundance of all shared expressed genes (FPKM ≥ 10: 4,190 genes for Ra infection, 4,194 genes for Rv infection) in the *Mtb* infected cells and their secreted exosomes. **(C)** RNA-binding proteins (RBP) recognition motifs of the *Mtb* infected cells and their released exosomes. Eight motifs are enriched in the exosomes (gcgcgss, ssagcgm, rygcgcb, cgcgc, ssgcgcs, crsmsgw, gckgghm, dgacrrr). **(D)** Boxplot showing the expressions of the genes with above-mentioned eight motifs in the *Mtb* infected macrophages and corresponding exosomes.

To further explore the transportation mechanism of these characteristic exosomal RNAs, we analyzed their RNA-binding proteins (RBP) recognition sites using the RBPmap database (Figure 5C), since previous reports have shown that RBPs contribute to the package of characteristic RNAs into exosomes [30]. The results revealed differential distribution of RBP recognition motifs between the *Mtb* infected cells and their secreted exosomes: most RBP recognition motifs were significantly enriched in the RNAs from *Mtb* infected cells, while eight were significantly enriched in the RNAs from secreted exosomes, all of which were associated with RNA binding, transport and translocation (Table S4) [27]. We then analyzed all the corresponding expressed genes containing these eight motifs. As expected, they showed much higher abundance in exosomes than those in cells (Figure 5D), suggesting selective secretion of some characteristic RNAs into exosomes through these eight RBPs. We further performed a GO analysis for these expressed genes in exosomes with these eight motifs, and interestingly, most of the enriched GO biological processes (4/6, ∼66.7%) were associated with cell adhesion (Figure S2). This suggests that these exosomal RNAs play roles in intercellular material transportation and signal transduction, as previously reported in exosomes [18, 19].

Further analysis indicated differential distribution for the eight exosome-enriched motifs between virulent and avirulent *Mtb* infections (Figure 5C), indicating their different transportation mechanism for the exosomal RNAs: two motifs (CRSMSGW and DGACRRR) showed a higher abundance in the exosomal RNA-transcripts derived from H37Ra infection, while five motifs (GCGCGSS, SSAGCGM, RYGCGCB, CGCGC, SSGCGCS) were significantly enriched in the exosomal RNA transcripts from H37Rv infection.

### Directed transport of *Mtb* RNAs to exosomes from macrophages

To explore the survival status of *Mtb*, we measured *Mtb* RNAs in infected macrophages and their secreted exosomes. As shown in Figure 6A, there was a much more significant enrichment of *Mtb* RNAs in exosomes compared to corresponding macrophages. Further analysis identified more *Mtb* genes in exosomes (3,114/3,840) than in corresponding macrophages (532/1,413) (Figure 6B). Among them, 2,631 (83.2%) and 2,436 (63.3%) *Mtb* genes were specifically detected in H37Rv- and H37Ra-derived exosomes, respectively, with only 49 (1.5%) and 9 (0.2%) identified in corresponding macrophages (Figure 6C). Furthermore, most co-expressed *Mtb* genes showed much higher abundance in exosomes than in corresponding macrophages: 96.01% (1,348/1,404) for H37Rv infections and 99.59% (481/483) for H37Ra infections (Figure 6C, E). Since the abundances of these co-expressed *Mtb* genes in exosomes was independent of those in cells (Figure 6D), their significant enrichment in exosomes could be attributed to their directed transport to exosomes from corresponding macrophages.

**Figure 6.**
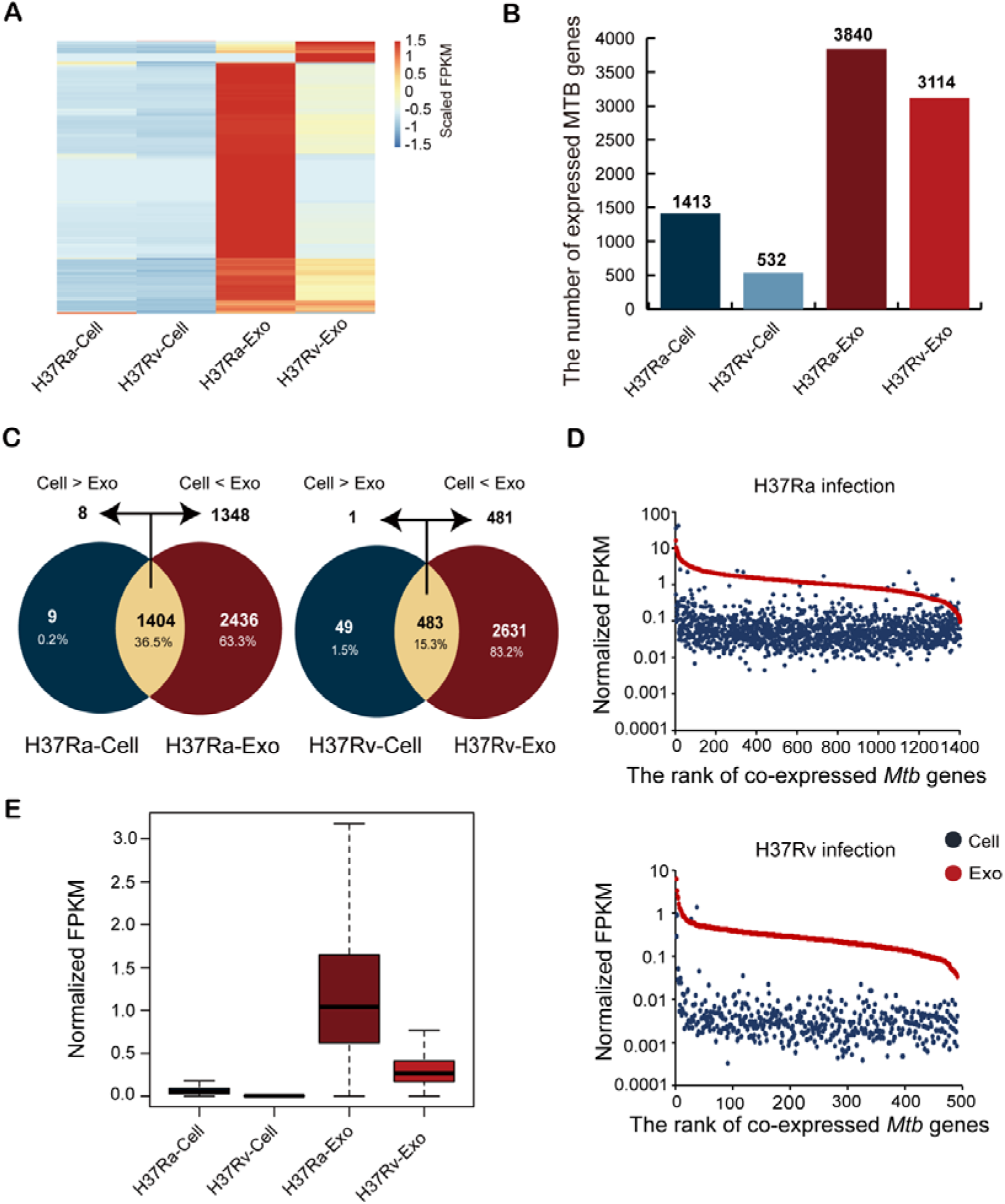
Expression profile of *Mtb* transcripts in the infected macrophages and secreted exosomes. **(A)** The abundance of *Mtb* transcripts among the infected macrophages and secreted exosomes. **(B)** The number of *Mtb* transcripts among the infected macrophages and secreted exosomes. **(C)** Venn plots showing the shared expressed *Mtb* genes in the infected cells and exosomes. **(D)** The abundance of all shared expressed *Mtb* genes (normalized FPKM values) in the infected cells and their secreted exosomes. The red and blue points represent the abundance of genes in exosomes and corresponding macrophages, respectively. Each point represents an *Mtb* gene. **(E)** Boxplot showing the abundance (normalized FPKM values) of *Mtb* genes in the infected cells and their secreted exosomes.

In addition, we observed a greater number of expressed genes and higher levels of *Mtb* in the exosomes derived from H37Ra-treated macrophages than in those from H37Rv-treated macrophages (Figure 6A, B, E). Consistently, previous study [12] has found that H37Ra can be eradicated through macrophage apoptosis under robust immune response, leading to secretion of *Mtb* RNA and other pathogen-associated molecular patterns (PAMPs) in the form of exosomes, while H37Rv could survive within macrophages by evolving an immune escape mechanism.

### High inhibitory effect of the treated exosomes and IFN-**γ** on *Mtb* infection

To further explore the inhibitory effect of treated exosomes and IFN-γ on *Mtb* infection, THP-1 differentiated macrophages were treated with exosomes derived from H37Rv or H37Ra infections in the absence or presence of IFN-γ for 24 h, which served as a vaccination method for the macrophages with the treated exosomes. After this, equimolar H37Rv was added, and the growth of cells and invading H37Rv were measured at 24 and 48 hours using MTT and CFU assays to evaluate the inhibitory effect (Figure 7A, B). The CFU assays showed that H37Ra-treated exosomes had a significantly higher antibacterial effect than H37Rv-treated exosomes and IFN-γ at 48h (p < 0.05), with a clear additive effect in the treated exosomes with exogenous IFN-γ added. Specifically, at 24 hours post H37Rv-infection, the inhibition started and the rates of “Macrophages + (H37Ra-treated exosomes) + IFN-γ”, “Macrophages + (H37Rv-treated exosomes) + IFN-γ”, “Macrophages + (H37Ra-treated exosomes)”, “Macrophages + (H37Rv-treated exosomes)” and “Macrophages + IFN-γ” for H37Rv were 62.0%, 55.5%, 49.2%, 32.6%, and 44.0%, respectively; after 48 hours, these values increased to 81.3%, 69.0%, 61.3%, 33.2% and 45.3%, respectively.

**Figure 7.**
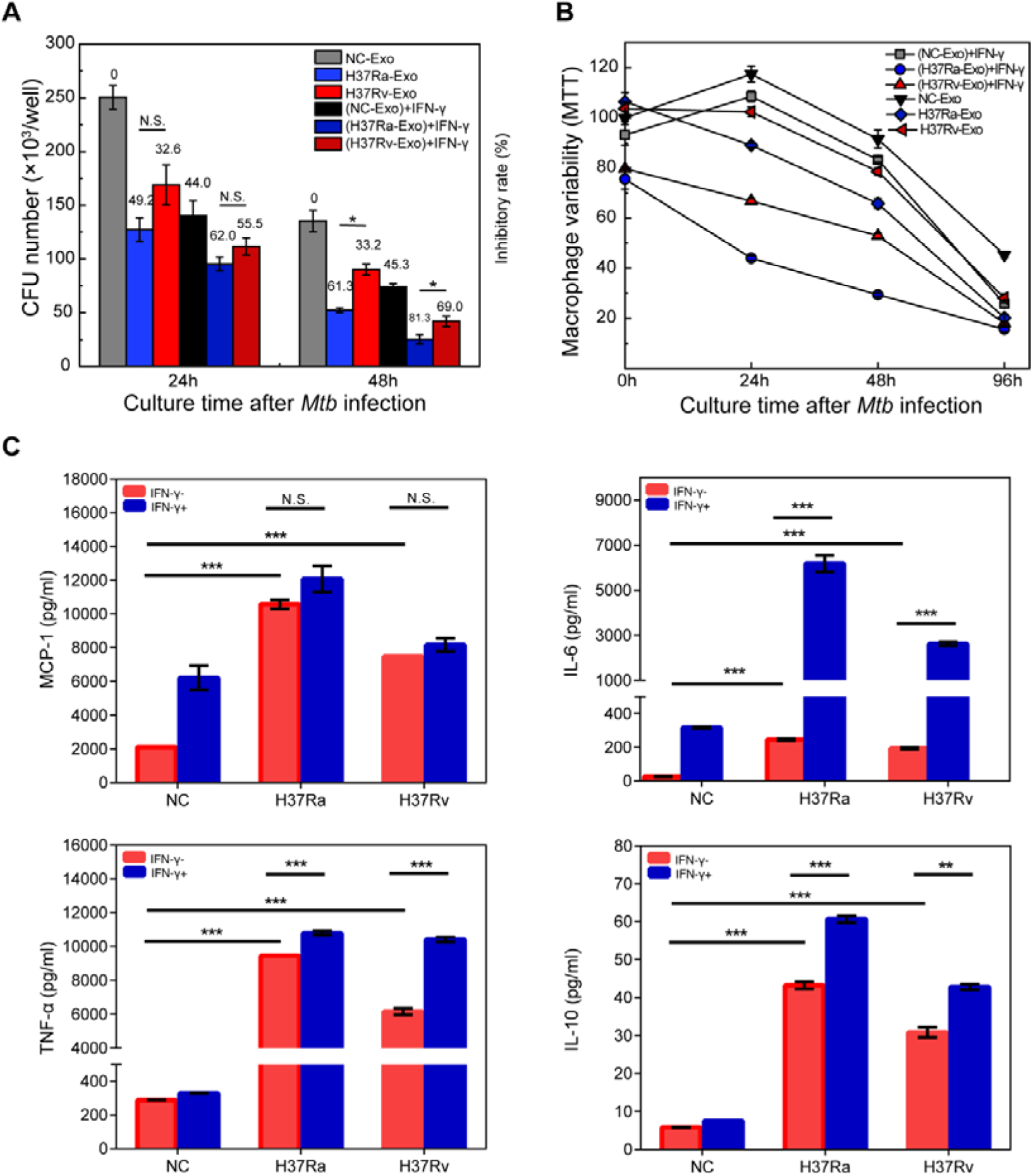
Anti-TB effect of exosomes from *Mtb* infected macrophages. **(A)** CFU assay showing the viability of the invading H37Rv in macrophages (MOI = 10) after exosome and/or IFN-γ treatments. **(B)** MTT assay showing the survival status of infected macrophages (MOI = 10) after exosome and/or IFN-γ treatments. **(C)** ELISA showing the expression of TNF-α, IL-6, IL-10 and MCP-1 in the macrophages treated with exosome and/or IFN-γ. Data are mean ± SEM and representative of three independent experiments. All the data are from three independent experiments with biological duplicates in each (mean ± SEM of n = 3 duplicates). N.S., not significant, *, p < 0.05, **, p < 0.01, ***, p < 0.001

On the other hand, our findings showed a negative correlation between the inhibitory effect of anti-H37Rv inhibitory effect (CFU assay) and cellular viability (MTT assay) (Figure 7A, B). In general, the ordering of cellular viability is as follows (from high to low) (Figure7B): “Macrophages (controls)” > “Macrophages + IFN-γ” > “Macrophages + (H37Rv-treated exosomes)” > “Macrophages + (H37Ra-treated exosomes)” > “Macrophages + (H37Rv-treated exosomes) + IFN-γ” > “Macrophages + (H37Ra-treated exosomes) + IFN-γ”. Here, the cellular viability of control and “Macrophages + IFN-γ” groups showed a growth trend during the first 24 hours of infection, followed by a significant decline after that time point [24]; the other groups showed a continuous decline from the beginning of the infection.

To investigate the mechanism underlying the inhibitory effect of the treated exosomes and IFN-γ on H37Rv infection, we measured the expression of several important cytokines in macrophages (IL-6, IL-10, TNF-α and MCP-1) (Figure 7C) since they have been reported to play important roles in the immune response to *Mtb* infection [31]. As expected, both “H37Ra and H37Rv-treated exosomes” could promote the expression of all the four cytokines in macrophages with “H37Ra-treated exosomes” stronger than “H37Ra-treated exosomes”, and the effect was enhanced after exogenous IFN-γ added to different extent.

### Differential enrichment of pro-inflammatory and immune-escape related *Mtb* proteins in “H37Ra-treated exosomes” and “H37Rv-treated exosomes”

To further explore why “H37Ra-treated exosomes” displayed higher antibacterial effect than “H37Rv-treated exosomes”, we performed comparative proteomic analyses of the *Mtb* treated exosomes. We identified 488 *Mtb-*derived differentially expressed proteins (DEPs) in the Ra/Rv group (ratio > 1.2, P-value < 0.05), and found that there were more DEPs in the H37Ra-treated exosomes (274/488, 56.15%) than in the H37Rv-treated exosomes (214/488, 43.85%). We then focused on 179 important functional proteins among the 488 DEPs that might be responsible for the antibacterial effect, including 65 antigens, 60 virulence factors, 95 membrane proteins, and 7 PE/PPE proteins. We found that there were more up-regulated DEPs in the H37Ra-treated exosomes than in the H37Rv-treated exosomes (Figure S3).

We then focused on the top 10 up-regulated DEPs in the H37Ra-treated exosomes, including six membrane proteins, two antigen proteins, one virulence factor, and one multi-functional protein (belonging to virulence factor, antigen protein, and membrane protein) (Table S5). Among them, eight proteins (MT1319, CTPA, Mmpl8, MT3772, Mmps3, RSMA, RPMI and RmlA) have been reported to be related to pro-inflammatory responses [32, 33]. In addition to these top 10 functional proteins, we also detected two enriched important immunogenic proteins in the H37Ra-treated exosomes: ESAT-6-like protein EsxS and 14 kDa antigen Hspx, which have been reported to stimulate a strong immune response against invading pathogens, including *Mtb* [34].

We also analyzed the top 10 up-regulated DEPs in the H37Rv-treated exosomes, including five membrane proteins, three antigen proteins, and two virulence factors (Table S5). Importantly, seven of these DEPs (Frdd, Devs, MftC, MmcO, AdhD, MT3488 and FadA) have been reported to play roles in *Mtb* survival in macrophages (pathogenicity) by inhibiting phagosome maturation [35–41], suggesting that H37Rv may escape immune surveillance by selectively secreting some specific proteins into exosomes.

## Discussion

### Selective/directed secretion mechanism of host and *Mtb* RNAs from the infected macrophages into exosomes

To the best of our knowledge, this is the first work to comprehensively analyze selective secretion and packaging of host and *Mtb* RNAs from the infected macrophages into exosomes. For host RNAs, our findings indicate that the selective packaging may be attributed to some RNA binding proteins (including RBM4, ZC3H10, RBM8A, FUS, PPRC1, SRSF1, and SAMD4A) with characteristic binding sites (such as gcfcgss, ssagcgm, rygcgcb, cgcgc, ssgcgcs, crsmsgw, and gckgghm). These findings are supported by previous studies: ribonucleoprotein A2B1 (hnRNPA2b1) has been reported to transport some characteristic miRNAs into exosomes by specific binding of GGAC motifs [30]; an RNA-binding protein YBX1 has been reported to transfer miR-233 into exosomes [42]; ELAVL1 has been documented to play roles in transporting some long non-coding RNAs into exosomes [43].

On the other hand, our findings showed much higher abundance of *Mtb* RNAs in exosomes than in cells (Figure 6A), which might be attributed to the directed transport of *Mtb* RNAs to exosomes from the phagolysosomes in macrophages [17]. In this process, pathogenic bacteria are phagocytosed into phagosomes, followed by the recruitment of lysosomes, which fused together to form phagolysosomes in the infected cells [17]. These infected pathogenic bacteria are then killed and degraded in the phagolysosomes, and the resulting exosomes are directly packaged and secreted. Therefore, the killing of bacteria in the phagolysosomal microenvironment may lead to significant enrichment of *Mtb* RNAs in exosomes compared to in cells.

### Macrophages vaccination with the treated exosomes containing some pro-inflammation *Mtb* proteins as promising TB subunit vaccine candidates

To investigate the underlying reason of the inhibitory effect of treated exosomes for microphages on *Mtb* infection (Figure 7A, B), we analyzed our proteomics data and found some enriched pro-inflammation *Mtb* proteins for immune protection in the exosomes (Figure S3). Herein, “H37Ra-treated exosomes” displayed a stronger antibacterial effect than “H37Rv-treated exosomes”, which could be due to some specially enriched immunogenic proteins in “H37Ra-treated exosomes”. Among the top 10 significantly up-regulated proteins in the H37Ra-treated exosomes (H37Ra/H37Rv) (Table S5), eight have been reported to play roles in pro-inflammation against *Mtb* infection: MmpS3 (top-1) is a member of MmpL/S family and a epitope/subunit vaccine candidate against *Mtb* infection; MmpL8 (top-5) is an important component of a promising vaccine candidate with significant protection against *Mtb* challenge in murine TB model; MT1319 (top-3) is an immune-reactive paratuberculosis antigen; CTPA (top-4) can induce innate immune responses in murine alveolar macrophages; RsmA (top-6) can block the transcription initiation of Esx secreted proteins to eliminate the infected *Mtb*; RpmI (top-8), a 50S ribosomal protein, can induce DC maturation and pro-inflammatory cytokine production (TNF-alpha, IL-1beta, and IL-6) through Toll-like receptor 4 (TLR4) recognition; RmlA (top-9) is related to PIM (phosphatidyl-myo-inositol mannosides) and can induce inflammatory response against *Mtb* infection. Overall, these proteins provide some promising candidates for TB subunit vaccines, and their therapeutic potential warrants further investigation.

Many studies have also demonstrated some enriched *Mtb* proteins as antigens in the exosomes against *Mtb* infection [35, 44]. For example, Bhatnagar *et al* found some *Mtb* antigen components in the exosomes isolated from bronchoalveolar lavage (BAL) fluid of TB patients [19]. They detected over 40 *Mtb* proteins and glycolipids in the exosomes secreted by infected macrophages, among which a 19 KDa lipoproteins was found to induce a pro-inflammatory response (such as TNF-α and IL-12 production) and the recruitment of macrophages and neutrophils [19].

Our findings also indicated a significant enrichment of *Mtb* immune escape related proteins in the H37Rv treated exosomes (Figure 4, Table S5), which have been reported to play roles in immune escape through inhibiting phagosome maturation and pathogenicity [36, 39, 45]. Herein, out of the top10 significantly up-regulated proteins in the H37Rv-treated exosomes, seven have been reported to favor their survival in cells: FrdD (top-3), a fumarate reductase, facilitates the survival of *Mtb in vivo*; DevS (top-5), as a part of a two-component system devR-devS, participates in the pathogenicity/intracellular survival of *Mtb*; MftC (top-6) is crucial in the synthesis of mycofactocin, which increases the survival fitness of *Mtb* within their specific niche; MmcO (top-7) protects *Mtb* from ROS attack; AdhD (top-8) plays important roles in the lipid biosynthesis for the formation of the mycobacterial cell envelope; MT3488 (top-9), a diterpene synthase, is essential for bacterial survival inside macrophages by inhibiting phagolysosome maturation and macrophage phagocytosis; FadA (top-10) contributes to *Mtb*’s survival, reactivation, and transmission by participating in the production of intracytoplasmic lipid droplets (LD).

Overall, the significant up-regulation of *Mtb* immune escape and *in vivo* survival related proteins in the H37Rv-treated exosomes sheds light on how *Mtb* evades host immunity for their survival *in vivo*. These proteins facilitate the signal transmission of immune-escape between cells for the survival of *Mtb in vivo*. Therefore, inhibiting these immune escape related proteins could disrupt the communication between cells and provide a new treatment strategy against *Mtb* infection.

### Simulative activation of interferon signaling pathways in macrophages even without IFN-**γ** expression

Interestingly, IFN-γ was predicted as the top upstream regulator in both H37Ra and H37Rv infected macrophages, and the “interferon signaling” pathway was highly activated. This is inconsistent with our traditional understanding about how IFN-γ performs functions during *Mtb* infection [29]. Typically, IFN-γ is produced by CD4+T and NK cells rather than macrophages, which plays a crucial role in macrophage activation for fighting against *Mtb* infection. Herein, the production of IFN-γ and other important antibacterial cytokines is a necessary part of the anti-*Mtb* cascading acquired immune responses *in vivo* (including PAMP recognition, antigen presentation, specific CD4+T cells activation, and antibacterial cytokines production) [9, 29]. Therefore, the significant activation of IFN-γ signaling pathways in macrophages, even in the absence of IFN-γ expression, is a surprising and interesting finding.

Both previous reports and our transcriptomic data indicated that IFN-γ was not produced in the infected macrophages (Table S3) [29]. However, the interferon signaling pathways were found to be significantly activated in the infected macrophages (Figure 3G, Figure 4A, 4B). So, how were these pathways activated? Our study has revealed a possible mechanism: by upregulating some downstream genes activated by other upstream genes, the interferon signaling pathways in macrophages can be simulatively activated to resist *Mtb* infection (Figure 4C, D). Therefore, we can infer that macrophages may be able to kill *Mtb* or other pathogenic bacteria in an IFN-γ independent, but a simulative way. Further studies are required to ascertain whether this compensatory strategy is truly present in the human body for fighting against *Mtb*. Moreover, the simulative activation of interferon signaling pathways in the absence of IFN-γ expression (Figure 4) suggests that it is a necessary pathway to kill invading *Mtb* in macrophages, and all roads lead to Rome.

### Important role of HMGB1 signaling pathway/*TNFRSF1B* in immune escape of virulent *Mtb* for *in vivo* survival

Our research has shown that H37Ra strain induces a stronger macrophage immune response than H37Rv strain in both phenotypes (Figure 2B, C) and genotypes (Figure 3), which was supported by one previous study on cytokine production in the H37Ra and H37Rv infected macrophages [46]. In our study, it is worth noting that only one immune related pathway, “HMGB1 signaling”, was significantly activated in the H37Rv-infected macrophage among all 10 activated immune response related pathways (Figure 4G), which has been reported to play a crucial role in immune escape for *in vivo* survival of *Mtb* by inducing necrosis [47]. Another study indicated that HMGB1 pathway promoted tumor immune escape by increasing myeloid-derived suppressor cells and IL-10 production [25]. Further, in our study, among four significantly upregulated genes enriched in the “HMGB1 signaling” pathway, *TNFRSF1B* has also been reported to inhibit apoptosis of infected macrophage by immune escape for *in vivo* survival of *Mtb* [48]. Collectively, based on our study and previous studies, we suggest that the weaker immune response in the H37Rv-infected macrophage may be mainly due to the significantly activated “HMGB1 signaling” pathway, which can manipulate cell death pathways (blocking apoptosis and activating necrosis) to escape immune response for further facilitating the survival of virulent *Mtb* in the infected macrophages. Therefore, inhibitors for “HMGB1 signaling” pathway as well as *TNFRSF1B* provides some promising therapeutic targets for the treatment of *Mtb* infection, and further studies on this is warranted.

### Limitation

While our study provides important insights into the macrophage immune response against *Mtb* infection, there are some limitations that need to be addressed. Our findings were obtained using an *in vitro* THP-1 cell model, and it is important to validate these results in more complex immune microenvironment. However, it is unfortunately that the validation experiments for some promising candidates can’t be performed, because the license for handling *Mtb* infection in animals in our hospital (Beijing Chest Hospital) was cancelled this year due to the unified planning of Beijing government. Despite these limitations, our study highlights the complexity of macrophage immune responses against *Mtb* infection, and further research is needed to fully understand the mechanisms involved in host-pathogen interactions.

## Conclusion

We all know that human immune responses in various immune cells decide the outcome of *Mtb* infection, macrophages play a crucial role in fighting against invading *Mtb*. However, a panoramic analysis of the immune mechanism in infected macrophages is still lacking. In this study, we conducted comprehensive analyses of the macrophages infected with avirulent and virulent *Mtb* (H37Ra & H37Rv) and dressed four concerns: 1. Avirulent *Mtb* stimulated robust immune-responses and apoptosis in macrophages to eliminate the invading *Mtb*; virulent *Mtb* induced severe necrosis and immune-escape for *in vivo* survival; 2. Macrophages kill *Mtb* in an IFN-γ independent but simulative way, highlighting the central role of IFN signaling pathway in anti-TB immune response; 3. We observed selective/directed secretion/transport of host and *Mtb* RNAs from macrophages to exosomes; 4. *Mtb* treated exosomes have great anti-TB effect which could be attributed to some pro-inflammation/immunogenic *Mtb* proteins. Overall, our findings revealed immune mechanism of macrophages in response to *Mtb* infection, and provided novel TB vaccine candidates.

## Materials and Methods

### Cell Line and Culture

Human THP-1 cells were cultured in 10ml RPMI 1640 medium (Biochrom, Germany) with 10% (v/v) FCS (Thermo Fisher Scientific, Germany), 1 mM L-glutamine, 100 U/ml penicillin and 100µg/ml streptomycin (Sigma-Aldrich, Germany). THP-1 monocytes were differentiated into macrophages by incubation with 50 nM Phorbol 12-myristate 13-acetate (PMA, Sigma Aldrich, USA) for 24 h.

### Bacterial Strains and Culture

Virulent and avirulent *Mtb* reference strain H37Rv and H37Ra were cultured at 37 °C in Middlebrook 7H9 medium (Difco) supplemented with 10% (vol/vol) oleic acid-albumin-dextrose-catalase (OADC; BD Biosciences), 0.5% glycerol, and 0.1% Tween-80 or on 7H10 plates (Difco) with 10% OADC and 0.5% glycerol.

### Cell Infection and *Mtb* staining

*Mtb* were inoculated into 7H9 medium and grown to OD = 1.0. The differentiated THP-1 macrophage cells (2 × 10^7^) were infected with *Mtb* at an MOI (Multiplicity of Infection) of 10. After 4-h phagocytosis, infected cells were washed with PBS before replacing with exosome-free RPMI 1640 medium. Cells and supernatant were collected after 24 hours post infection. Intracellular bacteria were identified using the Ziehl-Neelsen (ZN) stain according to manufacturer’s instructions and detected with microscopy.

### Exosome Extraction

Exosomes from *Mtb* infected THP-1 macrophages were isolated as previously described [22, 23]. Transmission electron microscopy (TEM) was employed to visualize exosomes using a JEM-1400 TEM (JEOL, Japan). The exosome size distribution was also measured using nanoparticle tracking analysis (NTA, Malvern, United Kingdom). Besides, TEM combined with immunogold labeling (surface marker CD9 and CD81) was employed to visualize exosomes (with an endoplasmic reticulum marker protein, Calnexin, as negative control) by using a JEM-1400 TEM (JEOL, Japan) (Figure S4).

### CFU and MTT Assay

The infected macrophages were washed with PBS, lysed in 0.5% Triton-X at 37 °C, serial diluted in PBS with 0.05% Tween-80, plated onto 7H10 plates, and counted for CFU after three weeks.

The infected macrophages were incubated with 10 μl MTT reagent at 8 h and 24 h post infection in cell incubator with 5% CO2 at 37°C for 4 h. The culture medium was then removed and 100 μl DMSO was added to each well. The absorbance was measured at OD_590_ using ELISA Reader (EnSpire^®^ Multimode Plate Reader).

### Cell Treatment with Exosome and/or IFN-γ

Differentiated THP-1 macrophages were treated with the extracted exosomes at ratio (exosome: cell) of 100:1, and/or IFN-γ (1000 U/ml) for 24hrs before infection.

### Cytokines Detection

Human TNF-α, IL-6, MCP-1 and IL-10 from the treated THP-1 macrophages were detected using MILLIPLEX^®^ milliplex assay (Millipore, Cat. HCYTOMAG-60K-08), and analyzed using Luminex200 platform.

### iTRAQ-LC-MS/MS analysis

Exosomes lysed in lysis buffer with protease inhibitors and concentrations measured using a BCA Protein Assay Kit (Thermo Scientific Pierce, USA). Samples (100 µg) from H37Ra and H37Rv-treated exosomes labelled with 8-iTRAQ reagents, acidified, mixed, and desalted with C18 tips as previously reported [49]. Eluted samples dried by SpeedVac for mass spectrometry.

LC-MS/MS was run on a Q-Exactive MS meter (Thermo Fisher Scientific, United States) equipped with Easy nLC (Thermo Fisher Scientific, United States) [50, 51]. Peptides loaded on trapping column and isolated with linear gradient of buffer B. Data-based top10 approach used to collect MS data in positive ion mode with dynamic exclusion duration set to 40 s. Resulting spectra analyzed with PD software and Mascot database. 1.2-fold change used as cut-off for biological significance based on normalized peptide ratios.

### RNA Sequencing, Data Processing and bioinformatic analysis

Total RNAs of the *Mtb* infected cells and isolated exosomes were extracted using RNAiso-Plus (Takara, Dalian, China) according to the manufacturer’s instructions. RNA concentration was measured using Qubit^®^ RNA Assay Kit in Qubit^®^ 2.0 Fluorometer. RNA integrity was assessed using the RNA Nano 6000 Assay Kit of the Agilent Bioanalyzer 2100 system (Agilent Technologies).

Sequencing libraries were constructed using NEBNext^®^ Ultra^TM^ Directional RNA Library Prep Kit for Illumina^®^. The library quality was assessed on the Agilent Bioanalyzer 2100 system. The RNA libraries were sequenced on the Illumina Hiseq 2500 Genome Analyzer platform in pair-end mode. Raw RNA-seq reads were filtered to obtain the clean RNA reads (FastQC) for subsequent analysis.

The clean data of each sample was mapped to the human reference genome GRCh38 using Tophat (version v2.1.0). The unique mapping reads were spliced and assembled using Cufflinks (version v2.2.1), and transcripts from the same gene were normalized to obtain the expression. Differentially expressed genes (DEGs) among samples were obtained using Cuffdiff, which were selected as follows: *P*-value < 0.05 and absolute fold-change (FC) ≥ 2.

To detect the *Mtb* transcripts, the clean data were mapped to the genomes of H37Ra and H37Rv using BWA. The expression levels of *Mtb* genes in cells and exosomes were obtained using Cufflinks.

We then performed bioinformatic analysis using the IPA (Ingenuity Pathway Analysis) software, which integrates knowledge manually curated by over 200 PhD experts. This software supports functional, pathway enrichment, and upstream regulator prediction analyses by considering the direction of gene expression changes based on biocurated literature evidence. The evaluation is performed using a Z-score, where an absolute Z-score greater than or equal to 2 is considered significant. A higher Z-score indicates a greater degree of activation of the related functions, pathways, or upstream regulators, while a lower Z-score suggests reduced activation. Detailed information about IPA and the Z-score can be found in Andreas et al. [52]. The Upstream Regulator Analysis in IPA represents a remarkable step forward in the field of transcription regulator prediction. Unlike other tools, IPA predicts which transcriptional regulators are involved and whether they are likely activated or inhibited based on biocurated evidence. IPA can then visualize this network of regulators and targets, explaining how the regulators interact with one another and their targets, thereby providing testable hypotheses for gene regulatory networks.

## Data availability

The raw sequencing data from this study have been deposited in the GSA-human database (https://ngdc.cncb.ac.cn/gsa-human), under the accession number: HRA004324.

## Code availability

The code for comparative transcriptomic analysis for macrophages infected with avirulent and virulent Mycobacterium tuberculosis has been submitted to BioCode at the National Genomics Data Center (NGDC), and are publicly accessible at https://ngdc.cncb.ac.cn/biocode/tool/BT7697.

## Acknowledgement

This work was supported by National Natural Science Foundation of China (82372260), Beijing Natural Science Foundation Haidian Original Innovation Joint Fund (L202023), and Funds for International Cooperation and Exchange of the National Natural Science Foundation of China (32061143024). We thank Prof Yongliang Zhao for the help with language.

## Supporting Information

Figure S1. RNA sequencing of H37Ra and H37Rv infected macrophages and released exosomes.

Figure S2. Bubble plot showing the enriched GO terms for the genes with above-mentioned eight motifs.

Figure S3. Bar plot representing 179 important functional differently expressed proteins (DEPs) in the H37Ra- and H37Rv-treated exosomes.

Figure S4. Identification of exosomes by TEM combined with immunogold labeling.

Table S1. The top 25 predicted upstream regulators for all DEGs in the H37Ra infected macrophages

Table S2. The top 25 predicted upstream regulators for all DEGs in the H37Rv infected macrophages

Table S3. Expression of Type I and II (IFN) interferons in the H37Ra/Rv infected Macrophages

Table S4. Specifically enriched RBP recognition motifs in the exosomes derived from H37Ra and H37Rv infections

Table S5. The top 10 significantly up-regulated differentially expressed functional proteins in the H37Ra- and H37Rv-treated exosomes

