## Supplementary data for "Comprehensive analysis of macrophages infected with avirulent and virulent *Mycobacterium tuberculosis* uncovers distinct immune mechanisms and anti-TB effect of treated exosomes"

**Supplementary Figures**

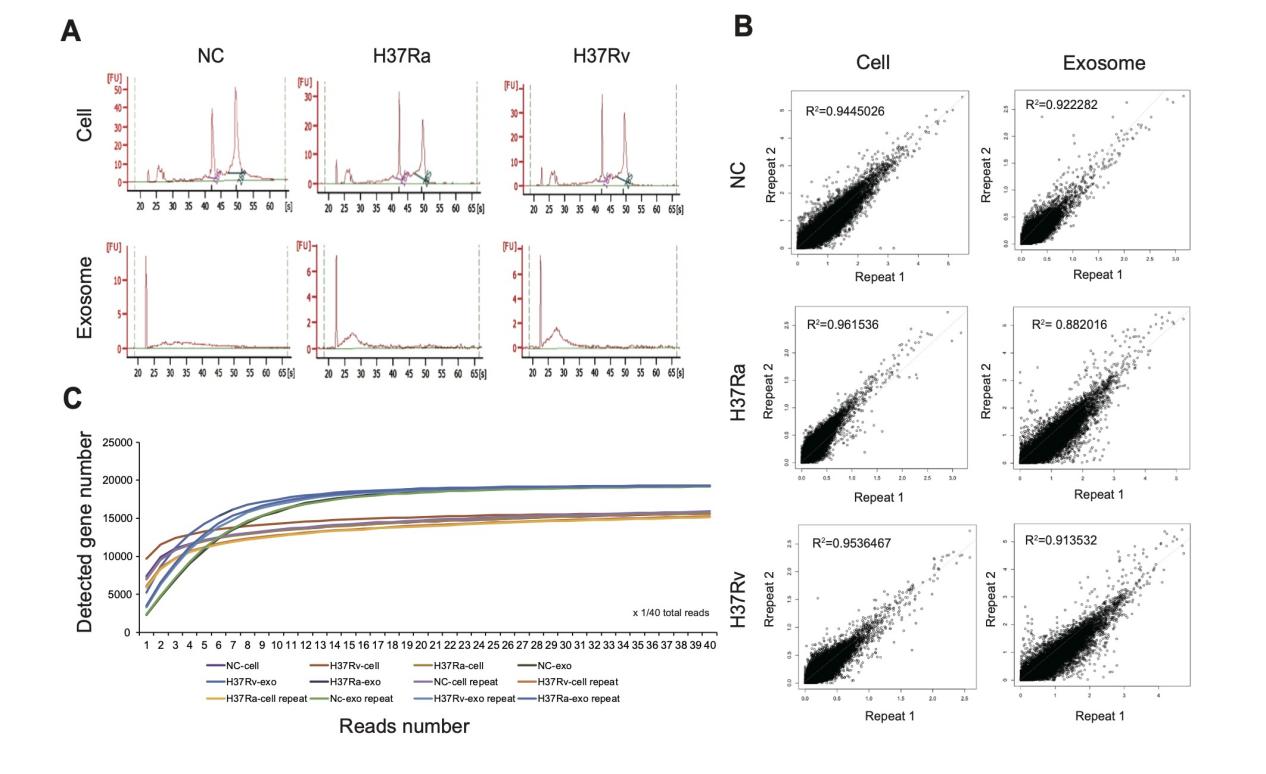

**Figure S1. RNA sequencing of H37Ra and H37Rv infected macrophages and released exosomes. (A)** Cellular and exosomal RNAs are assessed on the Agilent Bioanalyzer 2100 system (Agilent Technologies, Santa Clara, CA, United States). Typical 28S and18S rRNA peaks are detected in the cellular RNAs. Exosomal RNAs are mainly distributed between 22~32 nt. **(B)** Pointplot showing the correlation between two independent repeats. **(C)** Saturation curves of sequencing data.

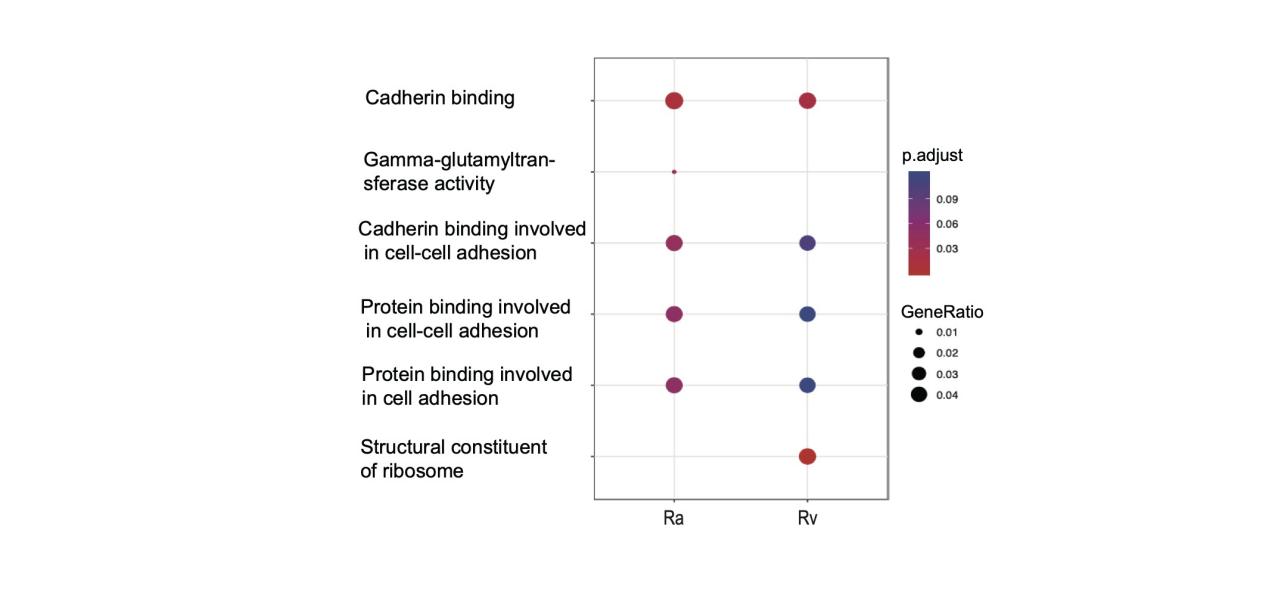

**Figure S2. Bubble plot showing the enriched GO terms for the genes with above-mentioned eight motifs.** The color indicates the adjust p value; the size of dots indicates the gene ratio.

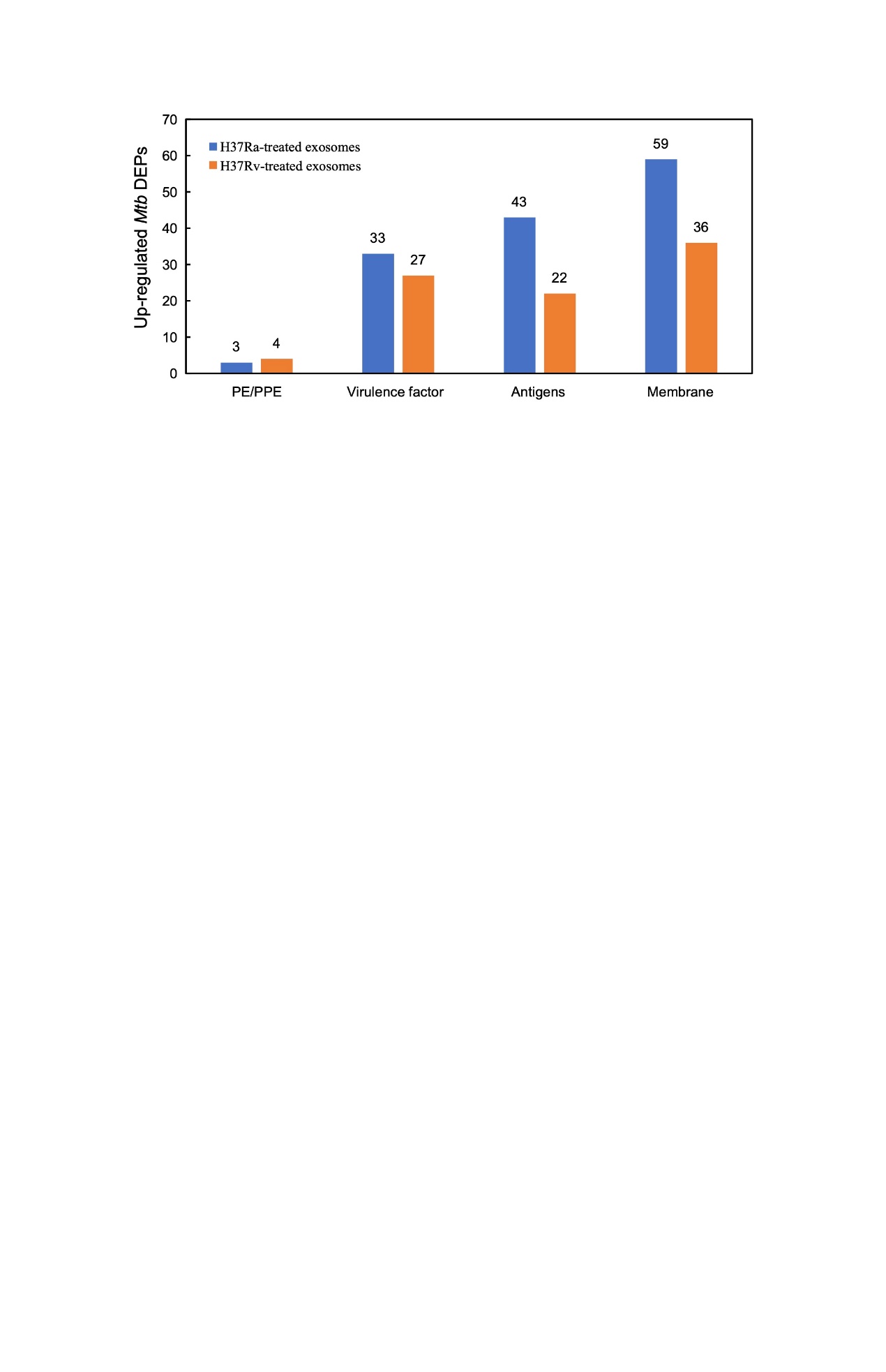

**Figure S3. Bar plot representing 179 important functional differently expressed proteins (DEPs) in the H37Ra- and H37Rv-treated exosomes.** The blue and orange bars represent the up-regulated Mtb DEPs in the H37Ra- and H37Rv-treated exosomes, respectively (Ra/Rv Ratio > 1.2, P-value < 0.05).

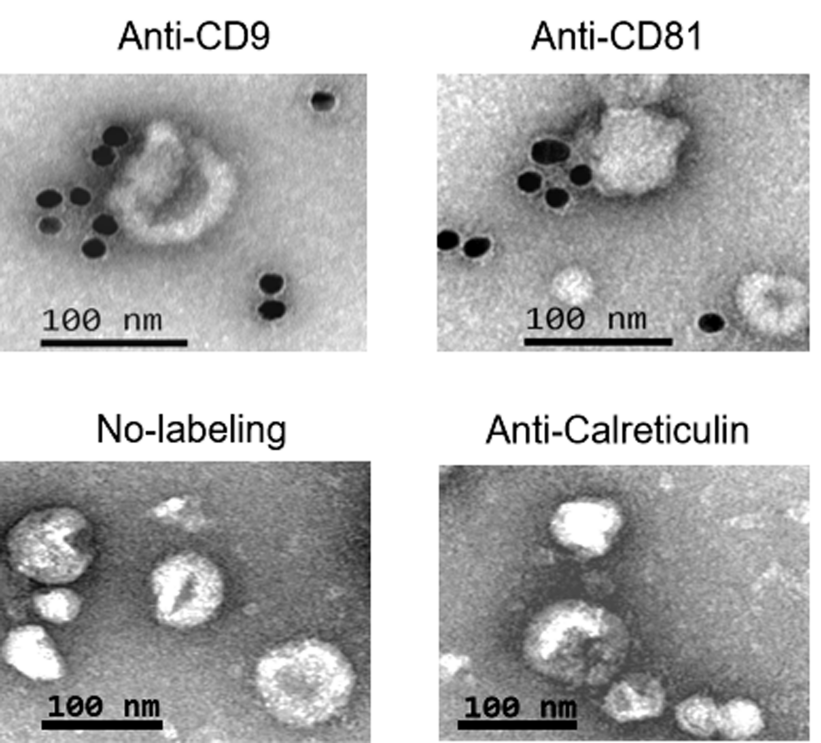

**Figure S4. Identification of exosomes by TEM combined with immunogold labeling.** Exosome surface marker CD9 and CD81 with rabbit anti-human antibodies of anti-CD9and anti-CD81 (SBI, United States), and an endoplasmic reticulum marker protein, Calnexin, as negative control was detected with anti-Calnexin antibody (Abcam, United States).

**Supplementary Tables**

**Table S1.** The top 25 predicted upstream regulators for all DEGs in the H37Ra infected macrophages

| **Upstream Regulator** | **Type** | **Z-score** | **P-value** |
| --- | --- | --- | --- |
| IFNG | cytokine | 6.064 | 7.53E-27 |
| poly rI:rC-RNA | biologic drug | 5.752 | 1.40E-32 |
| Interferon alpha | group | 5.244 | 2.67E-27 |
| Ifnar | group | 5.078 | 7.98E-31 |
| STAT1 | transcription regulator | 4.869 | 9.98E-28 |
| TLR3 | transmembrane receptor | 4.712 | 3.25E-31 |
| TNF | cytokine | 4.687 | 1.78E-18 |
| TLR9 | transmembrane receptor | 4.58 | 1.48E-22 |
| IRF7 | transcription regulator | 4.52 | 4.21E-23 |
| MYD88 | other | 4.454 | 8.08E-17 |
| IRF3 | transcription regulator | 4.342 | 6.48E-33 |
| IRF5 | transcription regulator | 4.323 | 2.90E-26 |
| IFNB1 | cytokine | 4.307 | 3.44E-26 |
| TICAM1 | other | 4.253 | 1.20E-20 |
| IL1B | cytokine | 4.171 | 5.57E-14 |
| TLR4 | transmembrane receptor | 4.121 | 1.36E-20 |
| Ifn | group | 4.028 | 9.15E-21 |
| IL21 | cytokine | 3.934 | 1.90E-12 |
| CHUK | kinase | 3.912 | 2.18E-14 |
| SAMSN1 | other | 3.9 | 1.75E-18 |
| IKBKB | kinase | 3.653 | 2.40E-10 |
| PTGER4 | g-protein coupled receptor | -3.889 | 7.87E-19 |
| IL10RA | transmembrane receptor | -4.359 | 1.16E-09 |
| TRIM24 | transcription regulator | -4.67 | 3.50E-22 |

**Table S2.** The top 25 predicted upstream regulators for all DEGs in the H37Rv infected macrophages

| **Upstream Regulator** | **Type** | **Z-score** | **P-value** |
| --- | --- | --- | --- |
| IFNG | cytokine | 5.698 | 3.33E-26 |
| poly rI:rC-RNA | biologic drug | 5.592 | 1.73E-29 |
| IRF3 | transcription regulator | 5.443 | 1.20E-37 |
| Ifnar | group | 5.28 | 5.47E-37 |
| STAT1 | transcription regulator | 5.054 | 1.40E-29 |
| IRF7 | transcription regulator | 5.028 | 1.01E-33 |
| Interferon alpha | group | 4.758 | 3.14E-23 |
| IRF5 | transcription regulator | 4.463 | 3.76E-30 |
| TLR3 | transmembrane receptor | 4.442 | 8.24E-21 |
| TICAM1 | other | 4.265 | 2.06E-17 |
| IFNB1 | cytokine | 4.256 | 2.52E-25 |
| NFATC2 | transcription regulator | 4.176 | 6.14E-16 |
| TLR9 | transmembrane receptor | 4.031 | 6.67E-14 |
| MYD88 | other | 3.946 | 1.23E-10 |
| Ifn | group | 3.903 | 2.19E-18 |
| SAMSN1 | other | 3.873 | 1.23E-14 |
| IL21 | cytokine | 3.873 | 1.06E-12 |
| TNF | cytokine | 3.846 | 4.93E-07 |
| TLR4 | transmembrane receptor | 3.581 | 4.45E-13 |
| TMEM173 | other | 3.44 | 4.94E-17 |
| ACKR2 | g-protein coupled receptor | -4.243 | 4.65E-27 |
| IL10RA | transmembrane receptor | -4.243 | 1.81E-10 |
| PTGER4 | g-protein coupled receptor | -4.408 | 2.99E-18 |
| TRIM24 | transcription regulator | -5.177 | 2.53E-32 |

**Table S3. Expression of Type I and II (IFN) interferons in the H37Ra/Rv infected Macrophages**

| **Gene name** | **Gene id** | **FPKM (H37Ra)** | **FPKM (H37Rv)** |
| --- | --- | --- | --- |
| IFNG | ENSG00000111537.4 | 0 | 0 |
| IFNA6 | ENSG00000120235.4 | 0 | 0 |
| IFNA8 | ENSG00000120242.3 | 0 | 0 |
| IFNA21 | ENSG00000137080.4 | 0 | 0 |
| IFNA5 | ENSG00000147873.5 | 0 | 0 |
| IFNA16 | ENSG00000147885.4 | 0 | 0 |
| IFNK | ENSG00000147896.3 | 0 | 0 |
| IFNB1 | ENSG00000171855.6 | 0.953749 | 0.390445 |
| IFNW1 | ENSG00000177047.6 | 0 | 0 |
| IFNL1 | ENSG00000182393.2 | 0.0775645 | 0.0736963 |
| IFNL2 | ENSG00000183709.7 | 0 | 0 |
| IFNE | ENSG00000184995.7 | 0 | 0 |
| IFNA10 | ENSG00000186803.3 | 0 | 0 |
| IFNA2 | ENSG00000188379.6 | 0 | 0 |
| IFNL3 | ENSG00000197110.8 | 0 | 0 |
| IFNA1 | ENSG00000197919.4 | 0 | 0 |
| IFNA7 | ENSG00000214042.1 | 0 | 0 |
| IFNWP18 | ENSG00000223684.1 | 0 | 0 |
| IFNA22P | ENSG00000224416.2 | 0 | 0 |
| IFNWP4 | ENSG00000225027.2 | 0 | 0 |
| IFNA20P | ENSG00000226393.1 | 0 | 0 |
| IFNWP9 | ENSG00000226597.1 | 0 | 0 |
| IFNA14 | ENSG00000228083.2 | 0 | 0 |
| IFNNP1 | ENSG00000230208.1 | 0 | 0 |
| IFNA11P | ENSG00000231195.1 | 0 | 0 |
| IFNWP5 | ENSG00000232138.1 | 0 | 0 |
| IFNWP15 | ENSG00000232281.2 | 0 | 0 |
| IFNA13 | ENSG00000233816.3 | 0 | 0 |
| IFNA17 | ENSG00000234829.3 | 0 | 0 |
| IFNA12P | ENSG00000235108.1 | 0 | 0 |
| IFNA4 | ENSG00000236637.2 | 0 | 0.040364 |
| IFNWP2 | ENSG00000237691.1 | 0 | 0 |
| IFNWP19 | ENSG00000238271.2 | 0 | 0 |
| IFNG-AS1 | ENSG00000255733.5 | 0.0814241 | 0 |
| IFNL3P1 | ENSG00000268510.1 | 0 | 0 |
| IFNL4P1 | ENSG00000272311.1 | 0 | 0 |
| IFNL4 | ENSG00000272395.5 | 0 | 0 |

**Table S4.** **Specifically enriched RBP recognition motifs in the exosomes derived from H37Ra and H37Rv infections**

| **motif** | **RNA-binding protein** | **Enrichment** | **Function** |
| --- | --- | --- | --- |
| gcgcgss | RBM4 | Ra & Rv | alternative splicing of pre-mRNA and translation regulation |
| ssagcgm | ZC3H10 | Ra & Rv | miRNA binding; mitochondrial physiology regulator |
| rygcgcb | RBM8A | Ra & Rv | spliced mRNAs |
| cgcgc | FUS | Ra & Rv | RNA transporting, pre-mRNA splicing and the export of fully processed mRNA to the cytoplasm |
| ssgcgcs | PPRC1 | Ra & Rv | transcription factor binding, nuclear receptor transcription coactivator activity |
| crsmsgw | SRSF1 | Ra & Rv | regulating alternative splicing |
| gckgghm | SAMD4A | Ra & Rv | mRNA binding; translation repressor activity |
| dgacrrr | FXR2 | Ra only | RNA binding, Translational Control and RNA transport. |

Note: Ra represents H37Ra infection group; Rv represents H37Rv infection group.

| **Table S5.** **The top 10 significantly up-regulated differentially expressed functional proteins in the H37Ra- and H37Rv-treated exosomes*** | | | | | | | | | |
| --- | --- | --- | --- | --- | --- | --- | --- | --- | --- |
| **Gene ID** | **Protein name** | **Ranking** | **Rv–Exo abundance** | **Ra–Exo Abundance** | **Ra–Exo/Rv–Exo Ratio** | **VF genes** | **Antigens** | **Membrane** | **References** |
| Rv2198c | MmpS3 | 1 | 27.6 | 172.4 | 6.25 |  |  | Y | PMID: 31652116 |
| Rv3219 | WhiB1 | 2 | 27.8 | 172.2 | 6.19 |  | Y |  | PMID: 24891105 |
| Rv1282c | MT1319 | 3 | 34 | 166 | 4.88 |  |  | Y | PMID: 18039835 |
| Rv0092 | CtpA | 4 | 39 | 161 | 4.13 |  |  | Y | PMID: 25703564 |
| Rv3823c | MmpL8 | 5 | 42.8 | 157.2 | 3.67 | Y | Y | Y | PMID: 29618733 |
| Rv3912 | RsmA | 6 | 43.1 | 156.9 | 3.64 |  |  | Y | PMID: 19951358 |
| Rv2578c | MT2655 | 7 | 43.7 | 156.3 | 3.58 |  |  | Y | PMID: 18083815 |
| Rv1642 | RpmI | 8 | 45.6 | 154.4 | 3.39 |  | Y |  | PMID: 25102137 |
| Rv0334 | RmlA | 9 | 49.2 | 150.8 | 3.07 | Y |  |  | PMID: 31728063 |
| Rv3671c | MT3772 | 10 | 49.4 | 150.6 | 3.05 |  |  | Y | PMID: 30366125 |
| Rv0859 | FadA | 10 | 155.7 | 44.3 | 0.28 |  |  | Y | PMID: 30644849 |
| Rv3378c | MT3488 | 9 | 157.1 | 42.9 | 0.27 |  | Y |  | PMID: 24475925 |
| Rv3086 | AdhD | 8 | 158.2 | 41.8 | 0.26 | Y |  |  | PMID: 11257547 |
| Rv0846c | MmcO | 7 | 158.6 | 41.4 | 0.26 |  |  | Y | PMID: 31278055 |
| Rv0693 | MftC | 6 | 159 | 41 | 0.26 |  | Y |  | PMID: 34311585 |
| Rv3132c | DevS | 5 | 162.1 | 37.9 | 0.23 | Y |  |  | PMID: 10970762 |
| Rv1368 | LprF | 4 | 166.8 | 33.2 | 0.2 |  |  | Y | PMID: 26201501 |
| Rv1555 | FrdD | 3 | 168.9 | 31.1 | 0.18 |  |  | Y | PMID: 18667562 |
| Rv1291c | MT1330 | 2 | 172.1 | 27.9 | 0.16 |  | Y |  |  |
| Rv0083 | MT0090 | 1 | 185.5 | 14.5 | 0.08 |  |  | Y |  |

* Rv–Exo abundance: the gene abundance in the H37Rv-treated exosome; Ra–Exo abundance: the gene abundance in the H37Ra-treated exosome; Ra–Exo/Rv–Exo Ratio: the ratio between the gene abundance in the H37Ra-treated exosome and that in the H37Rv-treated exosome.
